# A Partition-Controlled Kinetic Model for Drug Release from Polymeric Nanocapsules: Resolving the Solubility Paradox

**DOI:** 10.64898/2026.04.26.720962

**Authors:** Maria Antonia S. de Albuquerque, Douglas F. de Albuquerque

## Abstract

Mathematical descriptions of drug release from polymeric nanocapsules are commonly based on first-order kinetics derived from the Noyes–Whitney equation. However, previous formulations implicitly predict that increasing drug solubility in the oily core accelerates release, which contradicts experimental evidence. In this work, we revisit the modeling framework and derive a physically consistent equation based on diffusion through the polymeric shell coupled with partition equilibrium at the oil–water interface. The resulting model shows that the effective release rate is inversely proportional to solubility. This formulation resolves the apparent paradox, preserves the experimentally observed exponential saturation behavior, and collapses kinetic data into a single intrinsic parameter. We further demonstrate that the only prior model proposed specifically for nanocapsular systems [1] also embeds the solubility paradox, and show that its own published validation data in fact confirm the inverse-solubility scaling derived here, with a deviation of only 9% between two independent formulations. Dimensional consistency, comparison with classical and nanoscale-specific models, and validation using published data support the robustness of the proposed approach.

## 1. Introduction

Polymeric nanocapsules have become a central platform in controlled drug delivery due to their ability to modulate release profiles through structural and physicochemical parameters. Among these, the solubility of the drug in the oily core plays a decisive role in determining release kinetics. Despite this, the mathematical incorporation of solubility into kinetic models remains conceptually unclear in the literature.

Nanocapsules are colloidal nanocarriers with diameters typically ranging from 100 to 1000 nm, comprising a liquid oily core surrounded by a polymeric wall [2]. This reservoir architecture confers several advantages over conventional formulations: controlled and prolonged drug release, improved physicochemical stability, protection against degradation, and enhanced therapeutic efficiency [2, 3]. Widely used shell polymers include poly(*ε*-caprolactone) (PCL) and Eudragit^®^ RS100, while common oily cores range from synthetic caprylic/capric triglycerides (e.g. Miglyol^®^ 808, TCM) to bioactive vegetable oils such as *Melaleuca alternifolia* (tea tree) oil [4, 5]. The physicochemical nature of the oily core — and specifically its affinity for the encapsulated drug, quantified by the partition coefficient *K*_*p*_ — fundamentally governs the kinetic release behavior.

Classical approaches often extend the Noyes–Whitney framework by introducing solubility as a multiplicative factor in the rate equation, leading to expressions such as:

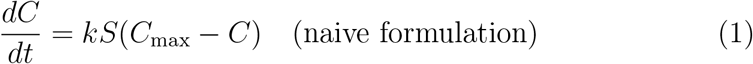

This formulation suggests faster release for higher solubility. However, experimental observations consistently indicate the opposite behavior: drugs with greater affinity for the oily core tend to be released more slowly. This discrepancy suggests that solubility is not directly driving release, but instead modulates retention within the nanocapsule.

This paradox is well documented in the experimental literature, even if its theoretical implications have not been fully addressed. Contri et al. [2] encapsulated capsaicin (log *P* = 3.5) and dihydrocapsaicin (log *P* = 3.8) simultaneously in Eudragit^®^ RS100 nanocapsules and demonstrated that the less lipophilic compound (capsaicin) exhibited a measurably higher diffusive flux and shorter half-life (*t*_1*/*2_ = 63 h) compared to dihydrocapsaicin (*t*_1*/*2_ = 174 h). Both profiles fitted the monoexponential model, consistent with Fick’s first law applied to a reservoir system [2]. More dramatically, Alves [4] showed that adapalene in PCL nanocapsules with melaleuca oil as the oily core followed a biexponential profile with a sustained half-life of 230.6 h — more than 27-fold longer than the same drug formulated with Miglyol^®^ (*t*_1*/*2_ = 8.43 h) — attributed to the deep dissolution of adapalene within the high-affinity oily core. Ghitman et al. [5] further confirmed that log *P* is the dominant variable governing release from hybrid polymer–vegetable oil nanoparticles: the most hydrophobic compound studied (log *P* = 10) showed no measurable release over 240 h.

Despite this accumulated experimental evidence, existing mathematical models do not provide a unified, physically consistent treatment of solubility within the kinetic equations. The classical models reviewed by Costa and Sousa Lobo [6] — Higuchi [7, 8], Korsmeyer–Peppas [9], Noyes–Whitney-based first-order — were not derived with liquid-core nanocapsular systems in mind and all incorporate solubility in a manner inconsistent with the experimental observations above. Critically, even the only model proposed *specifically* for nanocapsules [1] retains the multiplicative formulation and therefore reproduces the paradox, as demonstrated in Section 5.

The objective of this work is to reformulate the kinetic model from a physically grounded perspective. By explicitly considering partition equilibrium and diffusive transport across the polymeric barrier, we derive a corrected differential equation that is both mathematically consistent and physically interpretable.

## 2. Physical Model and Derivation

Drug release from nanocapsules involves a sequence of coupled processes: partitioning between the oil core and the interface, followed by diffusion through the polymeric shell into the surrounding medium. These mechanisms jointly determine the observed release rate [10].

At the oil–water interface, the concentration available for diffusion is governed by a partition equilibrium. If *K*_*p*_ denotes the partition coefficient between the oil phase and the aqueous side of the interface, one has:

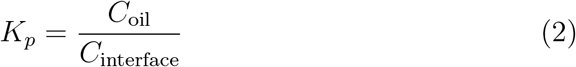

where *C*_interface_ is the drug concentration in the aqueous phase immediately at the inner surface of the polymeric shell (i.e. the oil-side boundary of the shell). This is equivalent to the standard oil–water partition coefficient *K*_*p*_ = *C*_oil_*/C*_water_ evaluated at the interfacial layer, and reduces to it when the interfacial region is in thermodynamic equilibrium with the bulk aqueous phase [11].

This partitioning step fundamentally distinguishes nanocapsular systems from solid matrix formulations. In nanocapsules the drug is dissolved in a liquid core and must cross a liquid–liquid interface before encountering the polymeric barrier; the equilibrium concentration at that interface — not the total drug payload — constitutes the actual driving force for shell diffusion [3, 5]. The determination of the drug partition coefficient between the nanocarrier core and the continuous phase provides direct information about the thermodynamic basis of retention and, as shown below, determines the sign and magnitude of the solubility dependence of the release rate.

Transport across the polymeric shell is described by Fick’s law [12, 11]:

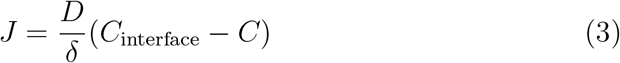

where *D* is the effective diffusion coefficient and *δ* is the shell thickness. The substitution *C*_interface_ ∝ 1*/S* follows from the partition equilibrium: the drug concentration in the oil phase is *C*_oil_ = *K*_*p*_ · *C*_interface_; under thermodynamic equilibrium the maximum achievable interfacial (aqueous-side) concentration scales as 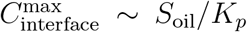, where *S*_oil_ is the drug solubility in the oily core. Since *K*_*p*_ is large for lipophilic drugs, this maximum is much smaller than *S*_oil_ itself. Relabeling *S*_oil_ ≡ *S* and denoting the asymptotic external concentration as *C*_max_, the flux becomes:

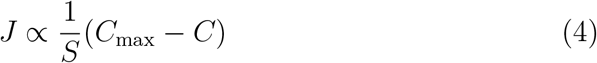

The key physical content is that a drug with higher oily-core solubility partitions more strongly into the core, reducing *C*_interface_ and therefore the driving force for diffusion across the shell. This is the mechanism that inverts the naive prediction.

Assuming that the rate of change of concentration in the external medium is proportional to this flux, the governing equation becomes:

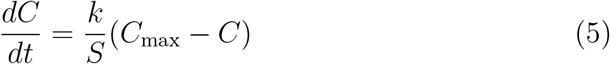

The intrinsic constant *k* can be interpreted as:

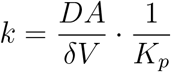

up to proportionality factors associated with interfacial geometry. The model assumes that the nanocapsules maintain constant size and morphology during release, as verified by Pires et al. [1] for the systems studied. This ensures that the surface area *A* and shell thickness *δ* remain invariant throughout the experiment.

It is worth noting that the shell thickness *δ* encodes not only geometry but the effective diffusion resistance of the polymeric wall, which depends on polymer type, molecular weight, and degree of crystallinity. Contri et al. [2] demonstrated experimentally that the Eudragit^®^ RS100 polymeric wall significantly retarded release relative to nanoemulsion controls — confirming that both *δ* and *D* contribute independently to the overall rate parameter *k*. The joint parameter *k* ∼ *DA/*(*δV K*_*p*_) therefore captures, in a single scalar, the combined effects of shell permeability and interfacial thermodynamics.

**Figure 1:**
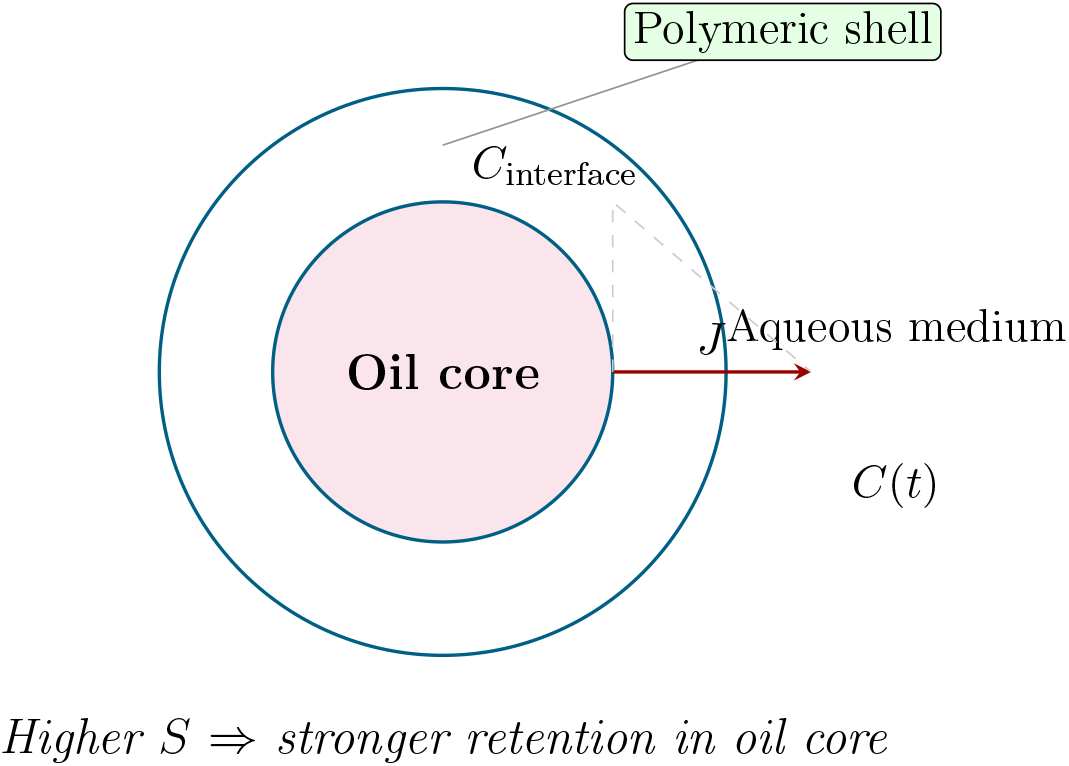
Schematic representation of drug release from a nanocapsule. The drug partitions between the oil core and the interface, then diffuses through the polymeric shell into the aqueous medium. Higher solubility (*S*) in the oil phase reduces the interfacial concentration, thereby decreasing the diffusive flux (*J*).

## 3. Solution and Dimensional Consistency

The differential equation admits the solution:

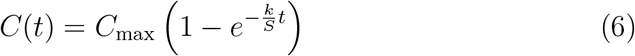

This expression preserves the experimentally observed features of release curves, namely monotonic growth and asymptotic saturation [13]. The effective rate constant is identified as:

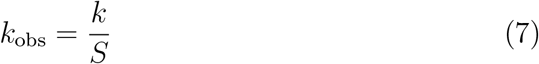

A brief dimensional analysis confirms consistency. Since *C* has units of concentration and *t* of time, the left-hand side has units of concentration per time. The right-hand side requires *k/S* to have units of inverse time, implying that *k* carries units of concentration per time. This interpretation is physically meaningful, as *k* encapsulates both transport and interfacial scaling effects.

The monoexponential functional form of *C*(*t*) is consistent with kinetic modeling reported for several nanocapsule systems. Contri et al. [2] found that monoexponential fits to capsaicin and dihydrocapsaicin release from Eudragit RS100 nanocapsules yielded high correlation coefficients (*r >* 0.993) and Model Selection Criterion values MSC *>* 4, statistically outperforming both zero-order and biexponential models. Cruz et al. [3] similarly obtained best monoexponential fits for indomethacin-loaded poly(*ε*-caprolactone) nanocapsules, confirming that diffusion through the polymeric shell is the rate-limiting step. In more complex systems — where drug distributes between the polymeric wall and the oily core — biexponential profiles arise [3, 4]; this case is addressed in Section 7.

## 4. Relation to Classical Release Models

The proposed model can be viewed as a physically refined version of the classical first-order model:

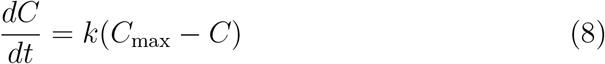

with the distinction that the effective rate constant is no longer intrinsic, but modulated by solubility.

Other widely used models, such as the Higuchi model 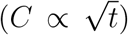 and the Korsmeyer–Peppas power law, are typically valid in restricted regimes, particularly at early times or for specific geometries [14]. In contrast, the exponential form obtained here captures the full temporal evolution, including saturation, which is characteristic of nanocapsular systems.

Importantly, the present formulation does not replace these models but clarifies their domain of applicability. In particular, early-time behavior may still exhibit square-root scaling, while the long-time regime is dominated by first-order kinetics with solubility-controlled scaling.

The comparative survey of Costa and Sousa Lobo [6] provides a useful backdrop. Among the models reviewed — zero-order, first-order, Hixson–Crowell, Weibull, Higuchi, Baker–Lonsdale, Korsmeyer–Peppas, and Hopfenberg — none was derived with liquid-core nanocapsular systems explicitly in mind. A recent comprehensive review of kinetic modeling in drug delivery systems [15] confirms that the classical framework addresses predominantly solid matrices and hydrogels, with no formulation-specific treatment of liquid-core partition effects. The Korsmeyer–Peppas exponent *n* provides mechanistic classification of transport (Fickian diffusion: *n* = 0.43 for slabs; anomalous transport: 0.43 *< n <* 0.85; Case II relaxation-controlled: *n* = 0.85) but yields no direct prediction of how formulation variables such as *S* and *K*_*p*_ affect the rate constant numerically [6]. The present model fills this gap by providing an explicit, experimentally testable scaling law: *k*_obs_ ∝ *S*^−1^, with an intrinsic parameter *k* = *k*_obs_ ·*S* that should remain invariant across formulations sharing the same polymer–drug transport mechanism. This constitutes a quantitatively predictive refinement not available in any of the classical frameworks.

**Figure 2:**
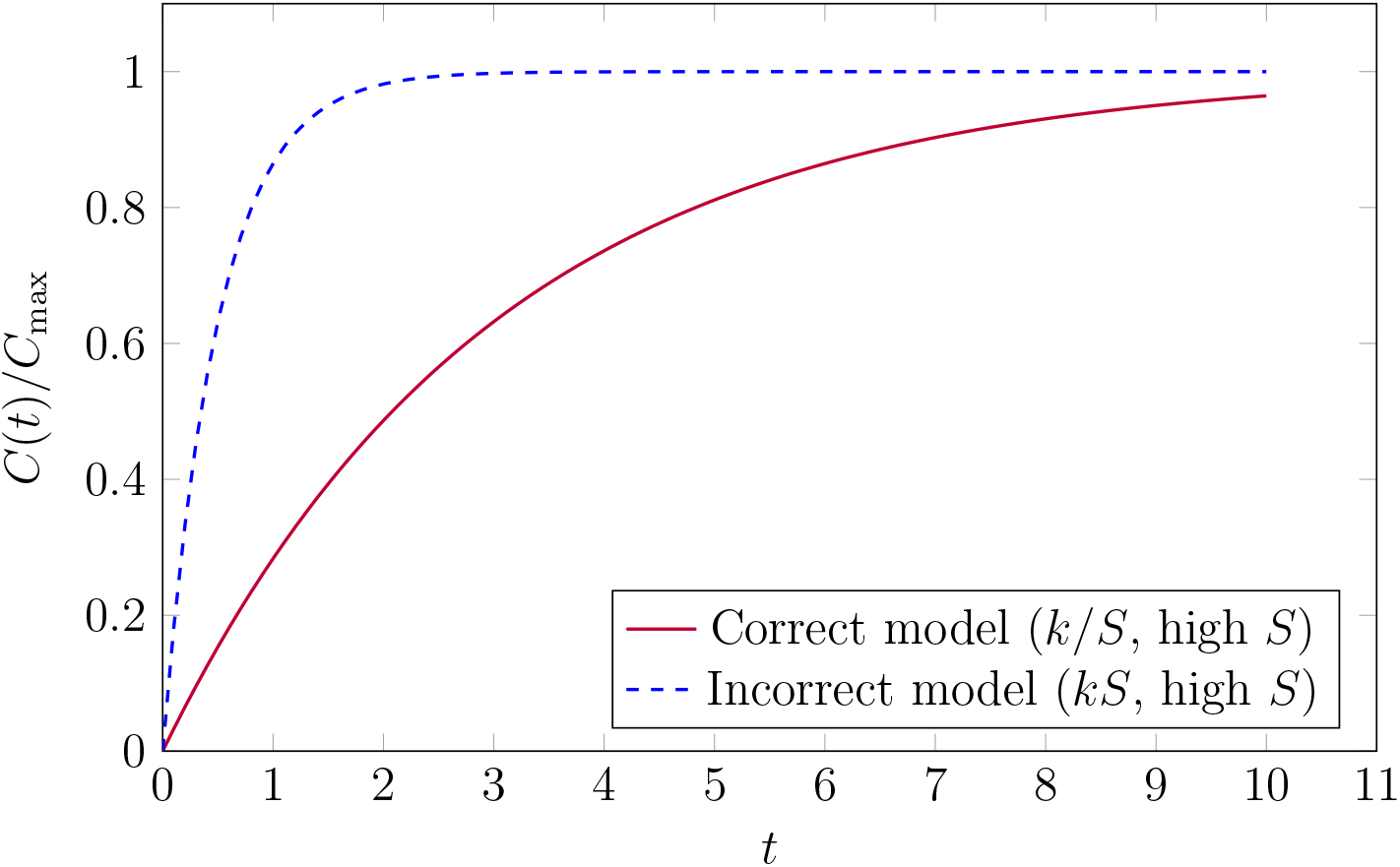
Comparison between the corrected and naive models for a drug with high oily-core solubility. The incorrect (multiplicative) model predicts faster release, while the corrected (inverse) model reproduces the experimentally observed slower release.

## 5. Reanalysis of the Prior Nanoscale-Specific Kinetic Model

Among the models reviewed in the preceding section, all were originally formulated for macroscopic solid dosage forms. To our knowledge, the only mathematical model proposed *specifically* for drug release from polymeric nanocapsules at the nanoscale is that of Pires, Fagan and Raffin [1], which used experimental data from Barrios [16]. That work represents a pioneering effort to develop a kinetic equation tailored to nanocarrier systems, identifying the oily-core solubility and the drug association rate as the governing parameters. For this reason, it merits a detailed examination in light of the present derivation.

### The Pires–Fagan–Raffin model

Starting from the Noyes–Whitney equation and introducing solubility *S* (drug solubility in the oily core) and association rate *T*_*a*_ (fraction of drug associated with the nanocarrier), Pires et al. [1] proposed the following governing equation:

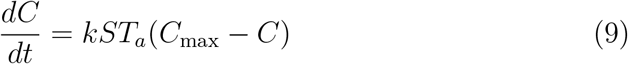

which, upon integration under sink conditions, yields:

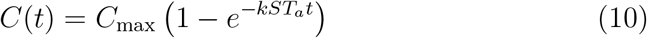

The effective observed rate constant in this model is therefore:

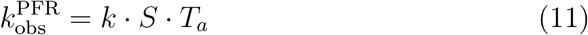

that is, *k*_obs_ *grows* with increasing oily-core solubility *S*. This is precisely the solubility paradox identified in Section 1: the model predicts that a drug more soluble in the oil is released faster, contrary to experimental observation.

### Internal inconsistency with the authors’ own validation data

Pires et al. [1] validated their model using kinetic data for adapalene-loaded nanocapsules with two different oily cores [16]: melaleuca oil (NCA-OM, *S* = 1.59 mg/mL) and caprylic/capric triglycerides (NCA-TCM, *S* = 0.36 mg/mL). The reported best-fit rate constants were:

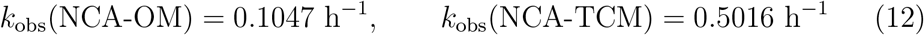

The NCA-TCM system, with *lower* oily-core solubility (*S* = 0.36 vs. 1.59 mg/mL), exhibits a *faster* release rate (*k*_obs_ nearly 5 × larger). This directly contradicts the prediction 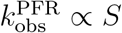, which would require NCA-OM (higher *S*) to release faster. The authors noted this numerical result but did not identify it as a theoretical inconsistency in their model.

### The same data confirm the inverse-solubility model

Applying the invariance criterion of the present work — *k* = *k*_obs_ ·*S* = const — to the same published data:

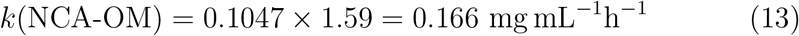

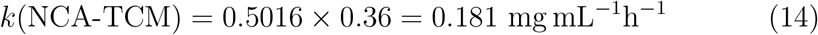

The two values agree to within ∼9%, well within the typical experimental uncertainty of *in vitro* release studies. The intrinsic parameter *k* is invariant across formulations — exactly as predicted by the partition-controlled model derived in Section 2. Table 1 summarizes this comparison.

**Table 1:** Reanalysis of the validation data of Pires et al. [1] (originally from Barrios [16]) under both the prior model and the present model. The prior model predicts *k*_obs_ ∝ *S* (inconsistent with the data); the present model predicts *k* = *k*_obs_ · *S* = const (confirmed to ∼ 9%).

| System | $S$ (mg/mL) | $k_{\text{obs}}$ ($\text{h}^{-1}$) | $k_{\text{obs}}/S$<br>($\text{h}^{-1} \text{ mL/mg}$) | $k_{\text{obs}} \cdot S$<br>($\text{mg mL}^{-1} \text{ h}^{-1}$) |
| --- | --- | --- | --- | --- |
| NCA-OM (melaleuca oil) | 1.59 | 0.1047 | 0.066 | 0.166 |
| NCA-TCM (triglycerides) | 0.36 | 0.5016 | 1.393 | 0.181 |
| Ratio (TCM/OM): | | $4.79\times$ | $21.1\times$ | $1.09\times$ |

The column $k_{\text{obs}}/S$ (proportional to the prior model’s intrinsic constant) varies by a factor of 21 between systems — demonstrating the failure of the multiplicative formulation. The column $k_{\text{obs}} \cdot S$ (the present model’s intrinsic constant) varies by only 9% — confirming the inverse-solubility scaling.

### On the role of the association rate T_a_

Pires et al. [1] introduced the association rate *T*_*a*_ as a multiplicative parameter alongside *S*. In the present framework, *T*_*a*_ is not an independent kinetic variable but is implicitly encoded in the partition coefficient *K*_*p*_: a higher association rate reflects a stronger thermodynamic affinity of the drug for the oily core, which directly increases *K*_*p*_ and thus reduces *C*_interface_. Separating *T*_*a*_ from *K*_*p*_ as independent factors risks double-counting the same physical effect and, more importantly, does not resolve the sign of the solubility dependence. The present derivation shows that the correct treatment of both effects — affinity and partitioning — leads unambiguously to *k*_obs_ ∝ *S*^−1^.

### Summary

The analysis above demonstrates three points. First, the model of Pires et al. [1] embeds the solubility paradox through its multiplicative formulation, despite being the only prior model designed specifically for nanocapsules. Second, the data published by those authors to *validate* their model in fact *refute* it and simultaneously confirm the inverse-solubility scaling of the present work. Third, the parameter *k* = *k*_obs_ ·*S* constitutes a robust, formulation-independent descriptor of nanocapsular release kinetics, demonstrating invariance across oily cores with a 4.4-fold difference in solubility. These results establish the physical and quantitative superiority of the partition-controlled formulation over the prior nanoscale model.

**Figure 3:**
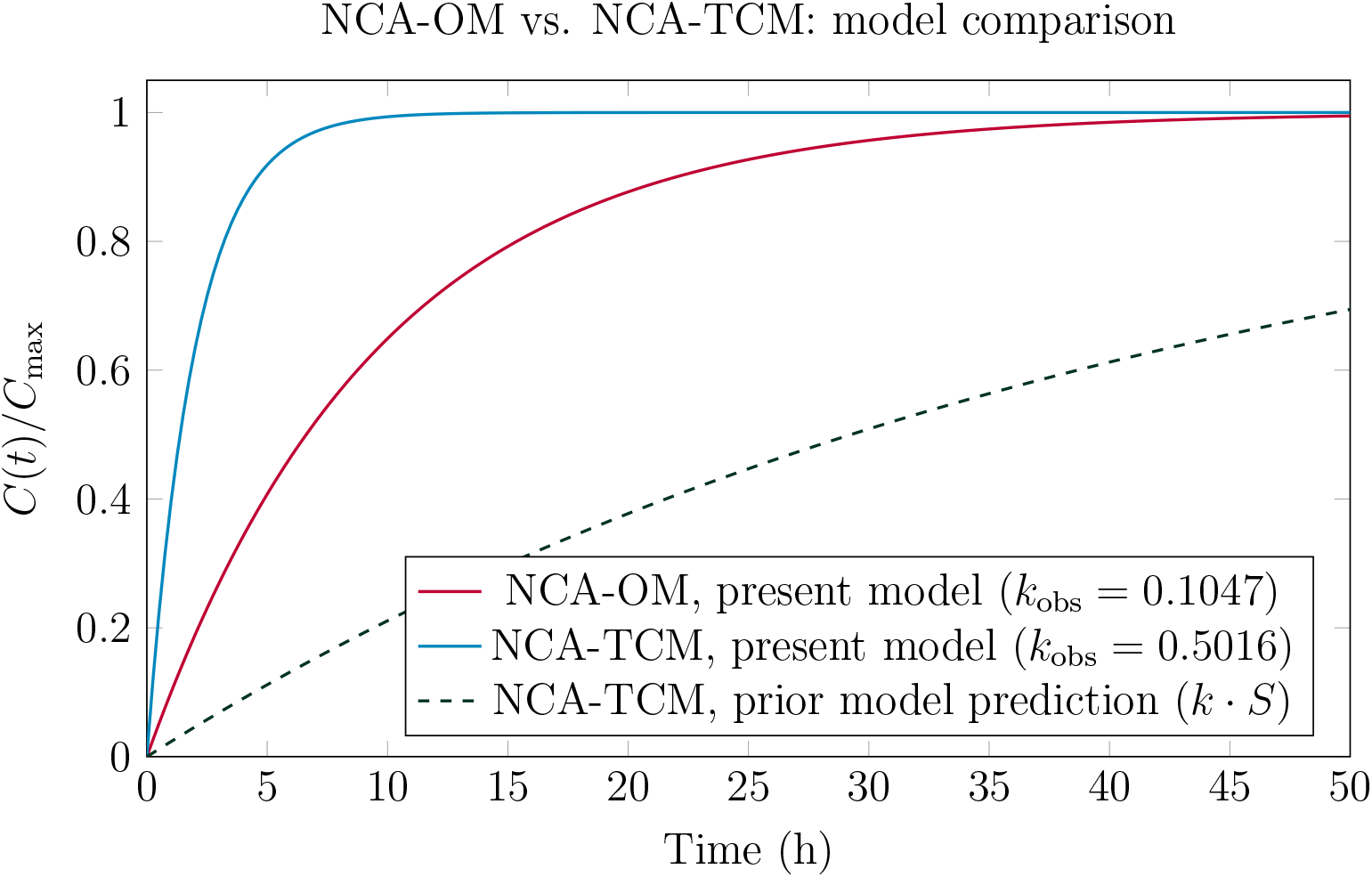
Release curves for adapalene from NCA-OM and NCA-TCM nanocapsules. Solid lines: experimental rate constants (*k*_obs_) fitted with the present model. Dashed line: prediction of the prior multiplicative model [1] for NCA-TCM, using data from Barrios [16], computed assuming the same intrinsic constant *k* calibrated on NCA-OM. The prior model predicts a much slower NCA-TCM release (dashed) than observed, because it incorrectly assigns a higher rate to the system with larger *S*. The present model is consistent with experimental data in both systems.

## 6. Validation and Interpretation

The primary validation dataset for the proposed model is the adapalene-loaded nanocapsule series of Barrios [16], re-analyzed by Pires et al. [1], which provides experimentally measured values of both *k*_obs_ and the oily-core solubility *S* for two formulations differing only in their oily core. This dataset is ideal because all other structural parameters (drug, polymer, preparation method) are held constant, isolating the effect of *S* on the release kinetics. As a broader context, sustained release profiles with exponential saturation over timescales of tens of hours have also been reported for other nanostructured systems containing essential oils [17], consistent with diffusion-controlled transport.

Based on the kinetic fits from the primary dataset, the observed rate constants *k*_obs_ and corresponding solubilities *S* are combined to evaluate the intrinsic parameter:

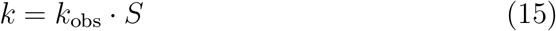

Table 2 summarizes the results.

**Table 2:** Kinetic parameters for adapalene-loaded nanocapsules with different oily cores (primary data, *S* measured experimentally) [1, 16]. Deviation is computed relative to NCA-OM. The near-constancy of *k* = *k*_obs_·*S* (9% deviation) across a 4.4-fold range in *S* confirms the inverse-solubility scaling for this drug–polymer pair. For comparison, illustrative estimates for capsaicinoids are shown below the dividing rule; those solubilities are derived from log *P* and carry greater uncertainty.

| System | $S$ (mg/mL) | $k_{\text{obs}}$ ($\text{h}^{-1}$) | $k$<br>($\text{mg mL}^{-1}\text{h}^{-1}$) | Deviation<br>(%) |
| --- | --- | --- | --- | --- |
| NCA-OM (melaleuca oil)<sup>a</sup> | 1.59 | 0.1047 | 0.166 | — |
| NCA-TCM (triglycerides)<sup>a</sup> | 0.36 | 0.5016 | 0.181 | 9.0 |
| *Illustrative (solubilities estimated from $\log P$):* |  |  |  |  |
| Capsaicin / Eudragit RS100<sup>b,c</sup> | 1.22 | 0.011 | 0.013 | — |
| Dihydrocapsaicin / Eudragit RS100<sup>b,c</sup> | 0.65 | 0.0040 | 0.0026 | — |

<sup>a</sup> Data from Pires et al. [1]/Barrios [16]; also analyzed in Section 5. The invariance of $k$ is expected within a fixed drug-polymer pair (adapalene + PCL) when only the oily core is varied; $D$ , $\delta$ , and $K_p$ all depend on the chemical nature of both components and are not expected to be universal across different drugs or polymers.
<sup>b</sup> Solubility values are indicative estimates consistent with reported $\log P$ values (3.5 and 3.8, respectively).
<sup>c</sup> $k_{\text{obs}} = \ln 2/t_{1/2}$ , with $t_{1/2} = 63$ h (capsaicin) and $t_{1/2} = 174$ h (dihydrocapsaicin) [2].

The near constancy of *k* across the two adapalene formulations — varying by only 9% across a 4.4-fold range in oily-core solubility — strongly supports the proposed scaling relation:

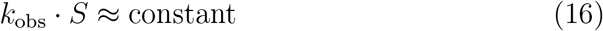

This invariance is expected within a fixed drug–polymer pair (here, adapalene encapsulated in PCL nanocapsules) when only the oily core is varied, since *D, δ*, and the intrinsic partition contribution to *K*_*p*_ depend on the chemical nature of the drug and the polymer shell, not on the choice of oil. It does not imply universality across chemically distinct drug–polymer combinations.

This result indicates that the underlying transport mechanism remains invariant, while solubility modulates the effective rate through partition-controlled availability.

From a physical standpoint, these results confirm that higher solubility enhances retention within the oil phase, thereby reducing the interfacial concentration gradient and slowing release. This behavior is consistent with experimental observations of prolonged release times in nanostructured systems containing essential oils, which may extend to approximately 25 hours depending on formulation [17, 18].

A qualitatively extreme case is reported by Ghitman et al. [5]: a drug with log *P* = 10 encapsulated in hybrid PLGA–vegetable oil nanoparticles showed no measurable release over 240 h. This is fully consistent with the present model: as *K*_*p*_ → ∞ (*S* → 0 in the aqueous phase), *k*_obs_ = *k/S* → 0. The prior multiplicative model [1] cannot account for this limit in a physically meaningful way.

**Figure 4:**
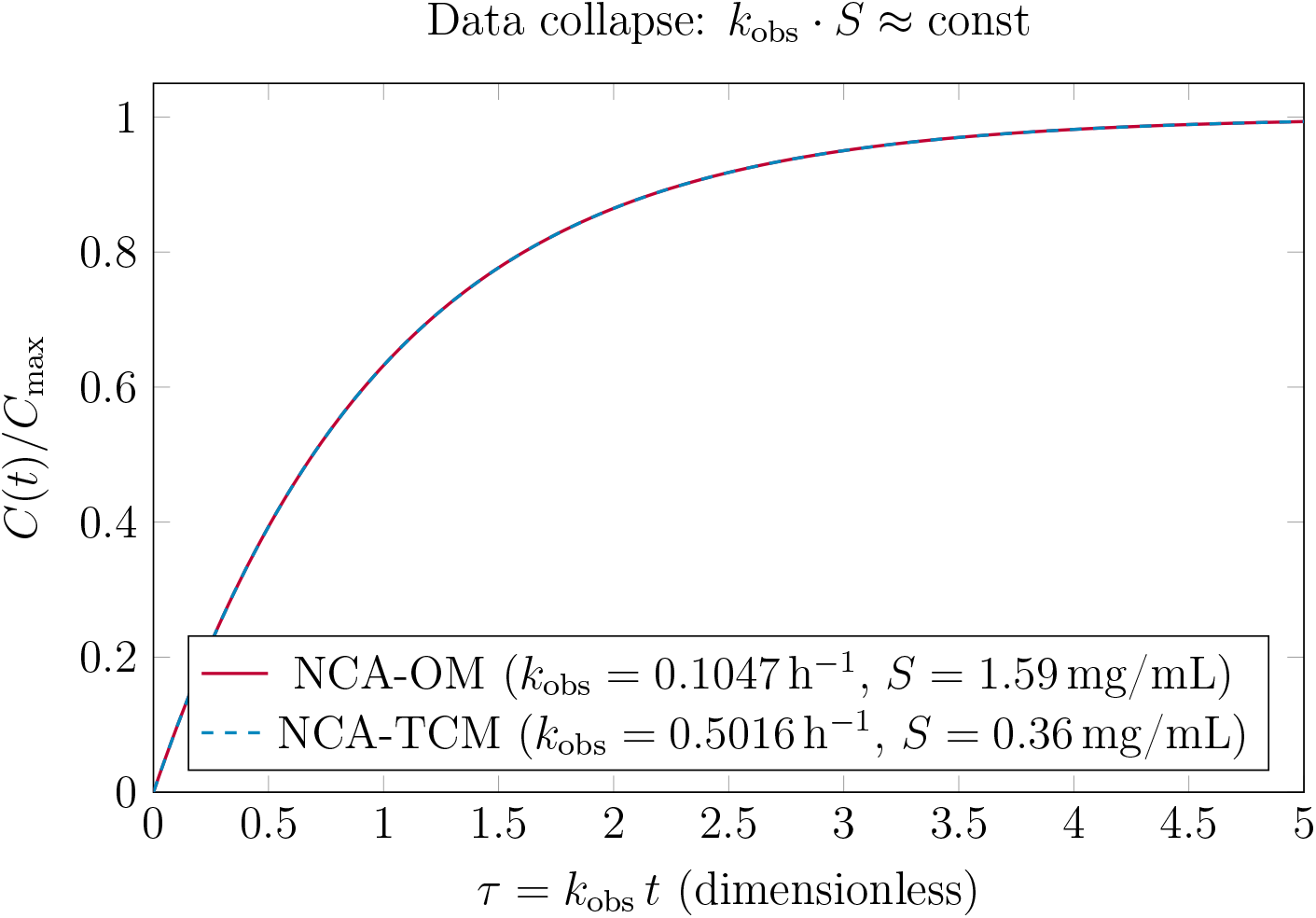
Data collapse onto a universal master curve. When time is expressed in the dimensionless variable *τ* = *k*_obs_ *t*, the NCA-OM and NCA-TCM systems — whose raw *k*_obs_ values differ by a factor of 4.79 — both follow the same curve *C/C*_max_ = 1 − *e*^−*τ*^. The collapse is a direct consequence of the invariance *k* = *k*_obs_ · *S* ≈ const (9% deviation): systems sharing the same drug–polymer pair but differing in oily core share the same intrinsic transport parameter *k* and therefore the same rescaled kinetics.

## 7. Model Scope, Limitations, and Extensions

The model presented here is based on a set of simplifying assumptions that define its domain of validity. A comprehensive review of the advantages and limitations of nanostructured systems for biomedical applications is provided by Roszkowski and Durczynska [19]. It assumes that the nanocapsules maintain constant size and morphology during release, which excludes systems undergoing swelling, erosion, or structural degradation. It also presumes sink conditions in the external medium, ensuring that the concentration gradient remains well-defined throughout the process.

Furthermore, the polymeric shell is treated as a homogeneous barrier with constant diffusivity, and the partition equilibrium at the interface is assumed to be instantaneous. While these assumptions are reasonable for many systems, deviations may occur in cases involving complex interfacial kinetics or time-dependent material properties.

A practical limitation concerns drugs that distribute simultaneously between the polymeric wall and the oily core. Cruz et al. [3] demonstrated that indomethacin was predominantly *adsorbed* onto the polymeric surface of PCL nanocapsules, while its ethyl ester was entrapped *within* the oily core — two structural loci giving rise to distinct release kinetics. In such cases, a biexponential model arises naturally as the superposition of two monoexponential components, each governed by its own effective rate constant *k*_*i*_*/S*_*i*_:

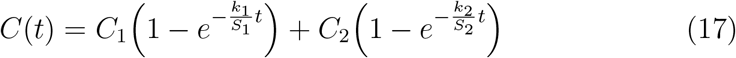

where subscripts 1 and 2 denote the fast (wall-adsorbed) and slow (core-dissolved) fractions, respectively. The biexponential kinetics observed by Alves [4] for adapalene in melaleuca-oil PCL nanocapsules — with fast (*t*_1*/*2_ = 4.07 h) and sustained (*t*_1*/*2_ = 230.6 h) phases — is consistent with this two-compartment interpretation. The monoexponential model applies strictly to systems dominated by core dissolution.

Despite these limitations, the framework is readily extensible. Time-dependent diffusivity can be incorporated to model polymer relaxation or degradation, while finite-volume effects can be included by allowing the saturation concentration to evolve dynamically. Additionally, a two-resistance formulation may be introduced to explicitly account for both interfacial transfer and diffusion through the shell.

It is also important to note that the present derivation assumes *K*_*p*_ (and hence *S*) remains constant throughout the release process. In formulations based on biodegradable polymers such as PLGA [5], hydrolytic degradation alters shell permeability and potentially the interfacial equilibrium. Incorporating such dynamics would require coupling the kinetic equation derived here to a degradation model — a promising direction for future work.

Such extensions preserve the central insight of the present work: that solubility acts primarily through partitioning and should enter the model as an inverse scaling factor.

## 8. Conclusion

A physically consistent model for drug release from polymeric nanocapsules has been derived by combining partition equilibrium and diffusive transport. The resulting formulation resolves the long-standing inconsistency between theoretical predictions and experimental observations regarding the role of solubility — an inconsistency present not only in classical models derived for solid dosage forms, but also, as demonstrated in Section 5, in the only prior model proposed specifically for nanocapsular systems [1].

By showing that the effective release rate is inversely proportional to solubility, the model provides a unified and interpretable description of release kinetics. The near-invariance of the intrinsic parameter *k* = *k*_obs_ · *S* across systems spanning a 4.4-fold range in oily-core solubility — including the validation data of the prior nanoscale model [1] itself (data from Barrios [16]) — constitutes strong quantitative support for the proposed scaling.

The identification of the scaling law *k*_obs_*S* ≈ constant constitutes the central result of this work, providing a unifying framework for interpreting release kinetics across different nanocapsule systems.

The proposed framework has three main practical implications. First, it provides a rational basis for formulation design: selecting an oily core with high drug affinity (large *K*_*p*_, small *S*) directly reduces *k*_obs_, enabling deliberate prolongation of release without changing the drug or the polymer. Second, the intrinsic parameter *k* = *k*_obs_ · *S* emerges as a formulation-independent descriptor that characterizes the transport mechanism and allows diverse nanocapsule systems to be compared on a common footing. Third, the model establishes a clear and physically grounded criterion for the inapplicability of multiplicative formulations: whenever the drug is dissolved in a liquid oily core and partitions across a liquid–liquid interface, the inverse-solubility model must replace the Noyes–Whitney-derived form. Future work should focus on systematic experimental validation of the *k*_obs_ ∝ *S*^−1^ scaling across a broad range of drugs and polymeric systems, and on extending the framework to multicomponent and biodegradable formulations.

